# Metabolic controls on carbon isotope fractionations during bacterial fermentation

**DOI:** 10.1101/2025.06.06.655161

**Authors:** Elliott P. Mueller, Verena B. Heuer, Jared R. Leadbetter, Kai-Uwe Hinrichs, Alex L. Sessions

## Abstract

Microbial fermentation facilitates the initial breakdown of organic matter into small molecules, and is thought to be the rate-limiting step for mineralization under anoxic conditions. Fermentation is understudied in modern and ancient biogeochemistry due to a lack of environmental biomarkers that would constrain its activity. It has long been assumed that fermentation, like respiration, does not express significant carbon isotope fractionations, precluding isotopic signals as a means of studying it in nature. Here, we tested this idea by growing pure cultures of four fermenting bacteria on glucose and measuring the carbon isotope compositions of the organic acids and alcohols produced. We found that fermentation exhibits a strong carbon isotope fractionation, ranging from -6‰ to +16‰, depending on the fermentation product. This range can even be observed within a single organism. Using bioisotopic models that track site-specific isotope enrichments through metabolic networks, we constrained the enzymes responsible for these fractionations. Our models reproduced in vivo organic acid *δ*^13^*C* values in all four organisms. These findings demonstrate that acetate ^13^*C*-enrichment is likely a consistent signature of fermentation. Furthermore, our study suggests that fermentation imposes an anaerobic trophic carbon isotope fractionation as organic carbon is passed from fermenters to secondary degraders like sulfate reducers. Looking to the geologic past, this trophic fractionation could have imprinted isotopic signals on the three billion year record of sedimentary organic carbon, specifically the inverse *δ*^13^*C* pattern of Precambrian acyclic isoprenoid and n-alkane biomarkers. Pervasive evidence of fermentation in the rock record suggests its under-appreciated role in biogeochemical cycles throughout Earth history.

**Significance Statement:** Microorganisms drive the global cycling of elements like carbon and regulate the Earth’s climate on both human and geologic timescales. Of particular importance is the microbial breakdown of organic matter, which generates greenhouse gases like methane. Fermenting bacteria play a crucial role in anaerobic carbon degradation. However, they remain largely invisible to our analyses. We explored the natural abundance of carbon stable isotopes in the molecular products of fermentation as a tool to study this metabolism in modern ecosystems and the ancient biosphere. We identified significant isotopic signals in molecules produced by fermenting bacteria and compared them to previously observed isotopic signals in molecular fossils in sedimentary rocks, which may provide evidence of microbial fermentation on the Precambrian Earth.

## Introduction

The fate of organic matter in anoxic environments exerts significant leverage on the Earth’s carbon cycle. The mechanisms and rates of organic degradation underpin the Earth’s long-term climate regulation, redox balance of the Earth’s surface, and nutrient cycling (i.e. through the burial of organic matter into ocean sediments) as well as short-term environmental responses to climate change (i.e. through greenhouse gas emissions from wetlands) [DeVries, 2022, Reay et al., 2018]. Fermentation is the first stage of anaerobic organic degradation during which biopolymers are broken down to small, labile molecules. Anaerobic microbial communities rely on these small molecules for carbon and energy, since sulfate reducers and many secondary degrading microbes cannot consume large biopolymers [Arndt et al., 2013, Arnosti, 2004, Beulig et al., 2018]. Fermentation products like acetate, formate, and ethanol have been observed in nearly every type of anoxic setting, highlighting the importance of this metabolism for carbon cycling. Acetate alone is the carbon source for as much as half of sulfate reduction and methanogenesis in the seafloor and two-thirds of methane production in terrestrial wetlands [Jørgensen et al., 2019, Conrad, 1999]. Fermentation is thus a major driver of the global carbon cycle. Here, we sought to address two fundamental questions about fermentation within the framework of carbon isotope biogeochemistry: 1.) Does fermentation impose predictable isotopic fingerprints onto its products? 2.) Might these fingerprints be used to study fermentation in modern environments and in the rock record?

There are several important differences between fermentation and respiration that underlie the structure of anaerobic carbon cycles. In contrast to respiring organisms, fermenting bacteria do not use electron transfer as their primary mode of energy conservation. Their largest energy source is substrate-level phosphorylation in central metabolism, which synthesizes ATP through the cleavage of phosphorylated compounds[White et al., 2012].While this metabolism is less efficient in terms of ATP generation per mole of substrate, it seems to enable fermenting organisms to break down larger molecules. The carbon substrates that fermenting bacteria can utilize are diverse, including amino acids, sugars, and biopolymers [White et al., 2012]. In contrast, at least in culture, anaerobically respiring bacteria and methanogens can only catabolize a small range of labile molecules. These physiological constraints on metabolism underpin the ecological structure of anaerobic microbial food webs in which fermenters provide carbon and energy from organic matter to the rest of the community.

Fermentation has a strong influence on carbon biogeochemistry. Most importantly, it may control the total rate of organic degradation and thus the rate of CO_2_ and methane generation in a given environment [Beulig et al., 2018]. Circumstantial evidence of this first appeared in the consistently low (*µ*M) abundance of fermentation products like acetate across different settings. Organic acid generation and consumption rates were balanced, indicating that hydrolysis and fermentation were the rate-limiting stage of organic matter degradation. Direct measurements of organic degradation rates throughout marine sediment profiles later confirmed this hypothesis [Arndt et al., 2013, Rothman, 2024]. In one study, the total rate of organic oxidation across the sulfate-methane transition zone (SMTZ) was unaffected by the shift in terminal consumption process from sulfate reduction to methanogenesis [Beulig et al., 2018]. Instead, the initial stages of organic matter decomposition set the pace of degradation throughout the sediment profiles. Fermenting microorganisms slowly released organic acids that were rapidly — and competitively — converted to CO_2_ and methane. These insights clearly demonstrate that fermentation can strongly influence rates of organic degradation, increasing the demand for tools to study these processes.

Despite the recognized importance of microbial fermentation, our ability to quantify fermentation in nature is limited. Open questions exist regarding the microbial species performing fermentation, the carbon substrates they use, and their metabolic activity. Oxidative substrates like sulfate and nitrate are routinely quantified in the environment as a means of tracing the metabolic activity of anaerobic respiration, but the organic substrates of fermentation are diverse and difficult to measure. The continuous turnover of fermentation products like acetate makes their abundances equally difficult to interpret. To our knowledge, no culture-independent, biological markers of fermentation have been developed either. Fermenting organisms have numerous pathways to disproportionate carbon and these pathways are phylogenetically widespread [Hackmann and Zhang, 2023]. As such, there are no characteristic enzymes or genes to screen for in environmental data. The ability to ferment cannot be identified via metagenomics, and currently requires confirmation via culturing. More recent work to classify fermenters based on genotype have had some success, but they are also biased toward cultured, fast-growing gut microbiota [Hackmann and Zhang, 2023]. Fermentation remains largely invisible to geochemical and biological tools in the environment.

Stable isotope fractionations between and within molecules have long been used to identify and even quantify metabolic activity. Indeed, the carbon isotope compositions of organic acids have been characterized in many anoxic environments and demonstrate demonstrably useful information. However, fermentation is assumed to have negligible isotopic fractionations during the production of these molecules (*<*3‰) [Heuer et al., 2010, 2006, 2009, Penning and Conrad, 2006, Conrad et al., 2021, Conrad and Claus, 2023]. Though previous studies demonstrated carbon isotope fractionations associated with fermentation, the enzymatic controls on these fractionations were not explored and they were assumed to have a negligible impact on the carbon isotope biogeochemistry of anoxic environments [Penning and Conrad, 2006]. Instead, the natural isotopic variability of fermentation products have been ascribed to consumption reactions, exchange with inorganic carbon or mixing with metabolic sources like autotrophic acetogenesis. [Lever et al., 2010, Franks et al., 2001]

Here, we asked whether fermentation has predictable isotopic fractionations that could be leveraged to study its activity in modern environments or in the geologic past. We studied four model bacteria (*Escherichia coli*, *Vibrio fischeri*, *Clostridium pasteurianum*, and *Zymomonas mobilis*) with well-constrained fermentation pathways. We chose these organisms because they represent a range of environments (enteric, marine, soil, and industrial, respectively), and they each ferment glucose as a sole carbon substrate using a distinct pathway, allowing us to sample some of the metabolic diversity that characterizes microbial fermentation. We hypothesized that the carbon isotope composition of the excreted products would reflect these different pathways. Indeed, the organisms expressed large and variable isotope fractionations between glucose and the fermentation products (+16‰ to -6‰). To connect the distinct metabolic pathways of each organism to their expressed isotope fractionations, we modeled each of their metabolic networks using a software known as Quantifying Isotopologue Reaction Networks (QIRN) [Mueller et al., 2022]. With these models, we identified the specific enzymes responsible for the observed fractionations, revealing the metabolic controls on the carbon isotope fractionations of fermentation. Finally, we discuss the implications of these isotope fractionations on the carbon isotope compositions of preserved molecular fossils in the Earth’s rock record, which may indicate the important role of fermentation throughout the Proterozoic Eon.

## Results

At the onset of this study, we hypothesized that microbial fermentation would express demonstrable and variable carbon isotope fractionations between the substrate consumed and the products excreted and that these fractionations would be primarily controlled by metabolic fluxes. To test this hypothesis, we grew four fermenters on the same substrate and measured the isotope composition of their excreted products. We then demonstrated that the observed fractionations were controlled by metabolic fluxes by creating bioisotopic models of each organism’s metabolism in QIRN. These models reproduced the broad range of isotope fractionations observed in the experiments across the four bacteria without changing the magnitude of kinetic isotope effects between organisms, confirming our original hypotheses. Below, our results are reported in more detail.

Under fermenting conditions, all four organisms consumed glucose and excreted organics acids and alcohols. Growth rates for *E. coli*, *V. fischeri*, *C. pasteurianum* are similar at 0.47, 0.434, and 0.473 hr*^−^*^1^. *Z. mobilis* had a slower (0.25 hr*^−^*^1^) growth rate (Figure S1). Two of the organisms (*E. coli* and *V. fischeri*) performed a mixed-acid fermentation, creating acetate, formate, ethanol, succinate, and lactate. *Z. mobilis* excreted exclusively ethanol, while *C. pasteurianum* made acetate, butyrate, and a minor amount of formate (Figures S2). By tracking the time-varying concentrations of all the organic products, their excretion rates were calculated (mM/gcell/hr) (Figure S3). Excretion rates were then normalized to the glucose uptake rate for each organism (mM/gcell/hr) to attain the zeroth-order fluxes found in Figure 1. These calculations were not possible for *E. coli* because time-varying samples were lost during transport. For *E. coli*, we assumed that the end-point substrate and product profile was representative of the uptake and excretion rates. From these data, a metabolic map of the metabolism was generated with flux-balance analysis (Figure 1). Under these assumptions and others regarding anabolism and reaction stoichiometry (see Methods), a unique metabolic network was determind for each organism studied. While similar in overall topology, the metabolic fluxes and branching ratios of each organism were different, even between *E. coli* and *V. fischeri* which have nominally the same mixed-acid fermentation pathway. Each organism excreted the expected organic products based on their known metabolic pathways. However, they all had different *δ*^13^*C* values (Figure 2A). This variability cannot be explained by changes in the substrate isotope composition, because the compound-specific *δ*^13^*C* value of glucose (-11‰) did not deviate by *>*1‰ in any of the cultures during growth (Table S1). The *δ*^13^*C* value of acetate was consistently enriched compared to glucose by 2-9‰. Ethanol and succinate were ^13^*C*-depleted (*δ*^13^*C* = -16 to -18‰) in *E. coli* and *V. fischeri*, more so even than palmitic acid which was made by all four organisms. In *E. coli*, *V. fischeri* and *C. pasteurianum*, palmitic acid was 4-5‰ ^13^*C*-depleted compared to glucose. Meanwhile, ethanol and palmitic acid from *Z. mobilis* were both more ^13^*C*-enriched than in other organisms. With a +5‰ *δ*^13^*C* value, formate created by *E. coli* had the largest isotopic fractionation from glucose. These results clearly demonstrate that fermentation expresses isotope fractionations (Figure 2A).

**Figure 1:**
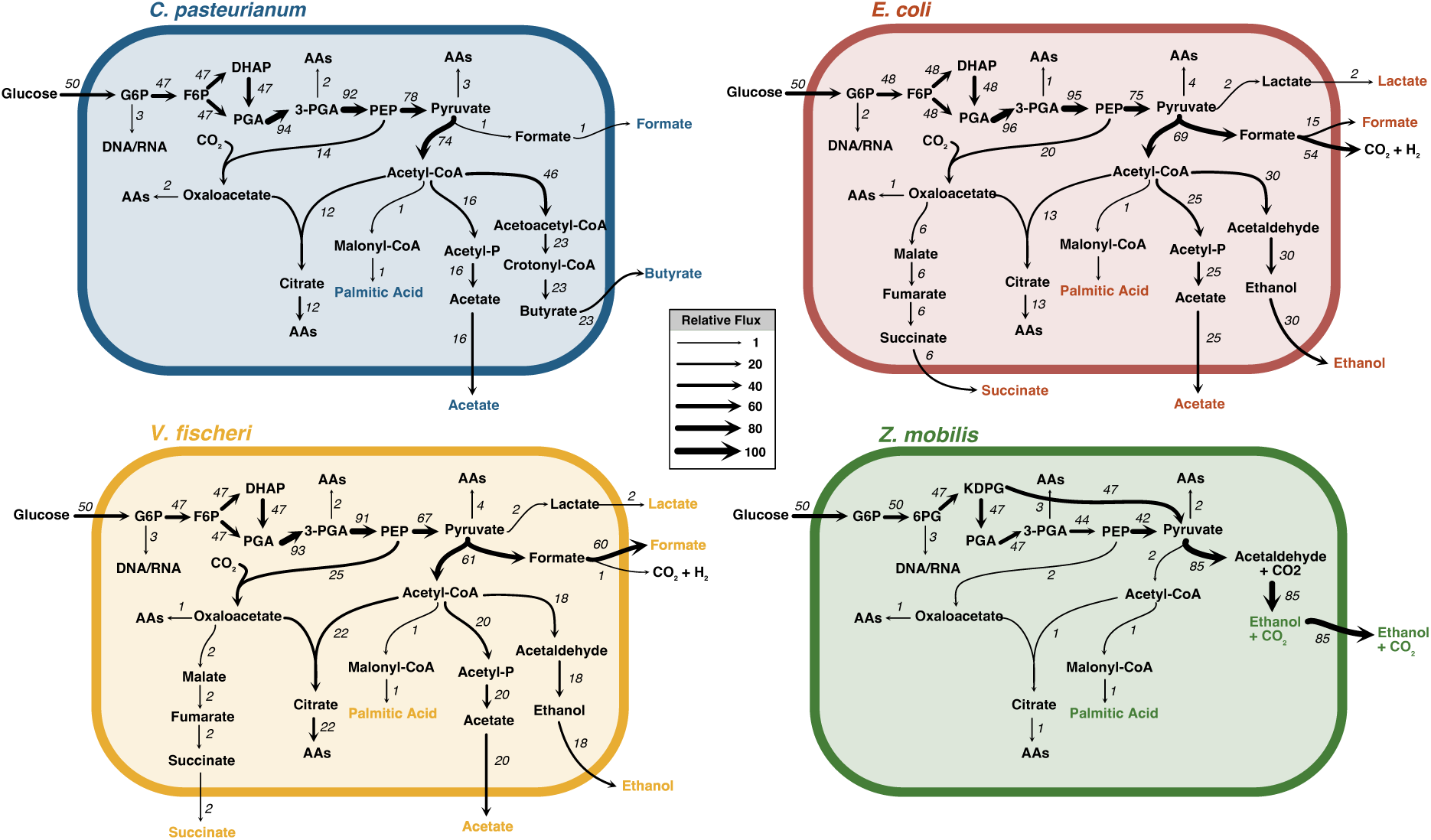
Metabolic flux maps generated in this study of all four organisms based on the quotient of the product excretion rates and glucose uptake rate normalized to 50. The flux terms represent moles of substrate through the reaction. Each model was implemented into QIRN with the measured intramolecular isotope composition of glucose and the compound-specific isotope composition of the products. All of the measured products are highlighted with colored font. Unique branching ratios at each metabolic node explain their variable net isotopic fractionations without needing to alter enzymatic isotope effects between organisms. Certain reaction pathways involving multiple steps are represented by a single reaction in the diagram for brevity. Abbreviations: G6P; glucose-6-phosphate, F6P; fructose-6-phosphate, DHAP; dihydroxyacetone phosphate, 3-PGA; 3-phosphoglycerate, PEP; phosphoenolpyruvate, AA; amino acids.

**Figure 2:**
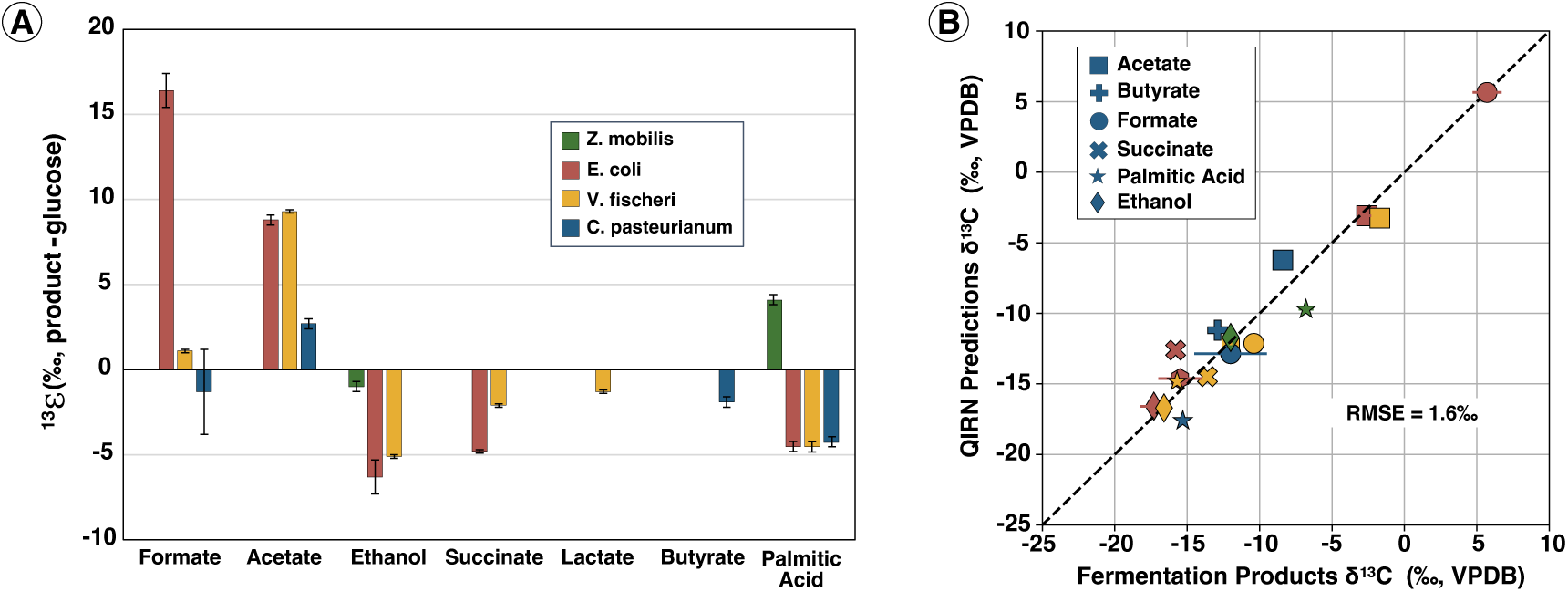
(A) Carbon isotope fractionation (^13^*ɛ*) between fermentation products and glucose observed in the four fermenting organisms grown in this study. Positive values indicate products with *δ*^13^*C* values more positive than the *δ*^13^*C* value of glucose. (B) Comparison of carbon isotope compositions of measured *in vivo* fermentation products results and model predictions. Dotted line represents one-to-one line. Error bars on both plots represents standard deviation of biological triplicates.

Our next goal was to understand the enzymatic reactions that were controlling this broad range of fractionations within and across organisms. To do this, we used a metabolic modeling tool called Quantifying Isotopologue Reaction Networks (QIRN). QIRN allowed us to model site-specific carbon isotope compositions of each metabolite in steady-state simulations of fermentation reaction networks (45-56 reactions and 48-60 metabolites). The models also incorporated the nonstochastic intramolecular *δ*^13^*C* heterogeneity found within the glucose molecules that these organisms consumed. For more information on how QIRN constructs and simulates reaction networks, see Supplementary Information and Mueller et al. [2022]. Model simulations of steady-state metabolic maps from Figure 1 were input into QIRN and site-specific ^13^C kinetic isotope effects (KIEs) were implemented into key enzymes within the fermentation pathways. With the fluxes determined, these KIEs became the free parameters of the models and were iteratively tested from 0.96 to 1.0 (^13^*k*/^12^*k*). For simplicity, secondary isotope effects were assumed to be negligible. The same enzymatic KIEs were applied to all four species. Measuring excretion rates does not constrain reaction reversibility, so all reactions were assumed to be irreversible, given that the cells were provided with abundant substrate to drive reactions in the forward direction. The model results were insensitive to KIEs for the majority of reactions (*>*80%), because they existed in unidirectional biosynthetic pathways (i.e. they had quantitative conversion of their substrate to product and could not express an isotope effect). Notably, model results were also insensitive to KIEs imparted by pyruvate decarboxylation and cleavage reactions (e.g. pyruvate dehydrogenase, pyruvate formate lyase), because pyruvate was nearly quantitatively (*>*90%) converted into a single product (e.g. acetyl-CoA) in every organism. The model results were more sensitive to the KIEs of enzymes at the formate, pyruvate, and acetyl-CoA nodes of metabolism, like those of formate dehydrogenase (FDH), acetyl-CoA carboxylase (ACC), citrate synthase (CS), alcohol dehydrogenase (ADH), and phospho-acetyl transferase (PTA) (Table 1, Supplementary Figure S4). The sensitivity of these enzymes was a consequence of the measured data used to fit the model. The molecules measured in this study had precursors at the nodes of metabolism influenced by these enzymes, so the KIEs of these enzymes had a strong influence on model results. Other enzymes could be expressing isotope effects (e.g. to synthesize amino acids), but we do not have measured constraints on those nodes of metabolism and they do not influence the isotope composition of the measured products so we assume they are unity here. The optimized KIEs found are in Table 1. Model outputs matched the data equally well (*<*3‰ root-mean square error, RMSE) within a range of 10‰ around most of the KIE values listed in Table 1. The chosen optimized KIEs in Table 1 fit the data (n = 17) with a RMSE of *<*1.6‰ (Figure 2B). The optimized fit was possible even though we forced all four organisms to express the same enzymatic KIEs. The only difference in the models is the network topology. The ability to fit data across these four diverse organisms using the same set of isotope effects suggests that enzymatic KIEs between organisms are similar.

**Table 1:** Best-fit KIEs for key enzymes in fermentative metabolisms.

| Enzyme | Reaction | Site | KIE |
| --- | --- | --- | --- |
| ADH | acetyl-CoA → acetaldehyde | C-1 | $0.973 \pm 0.005$ |
| ACC | acetyl-CoA + CO<sub>2</sub> → malonyl-CoA | C-2 | $0.973 \pm 0.005$ |
| ACAT | 2 x acetyl-CoA → acetoacetyl-CoA | C-2 | $0.987 \pm 0.005$ |
| | | C-3 | $0.987 \pm 0.005$ |
| FDH | formate → CO<sub>2</sub> | C-1 | $0.977 \pm 0.005$ |
| CS | oxaloacetate + acetyl-CoA → citrate | C-6 | $0.98 \pm 0.01$ |
| PTA | acetyl-CoA → acetate | C-1 | 1.000 |

**Abbreviations:** Alcohol Dehydrogenase (ADH), Acetyl-CoA Carboxylase (ACC), AcetylCoA Acyltransferase (ACAT), Formate Dehydrogenase (FDH), Citrate Synthase (CS), Phosphate acetyltransferase (PTA)

Overall, our data and models indicate that fermentation preferentially partitions ^13^C into excreted products and ^12^C into carbon dioxide and biomass (Figure 4). Over half of the carbon consumed by these fermenting bacteria was excreted as small molecules, and this pool was on average 3‰ enriched relative to glucose. The remaining carbon was either lost as carbon dioxide or used to build biomass. Simple mass balance predicts that both of these pools were ^13^*C*-depleted. Indeed, all of the biomass carbon isotope compositions were depleted (-11 to -17‰) providing a notable exception to the rule of thumb “you are what you eat +1‰”. While inorganic carbon was not measured in these studies, we calculated that the CO_2_ produced by these cultures would only need to be 1-2‰ lower than the substrate to attain isotopic mass balance, assuming carbon was only transformed into organic acids, biomass, and CO_2_ and biomass accounted for 10% of the consumed carbon, though the calculated CO_2_ *δ*^13^*C* value is not sensitive to biomass contribution.

**Figure 3:**
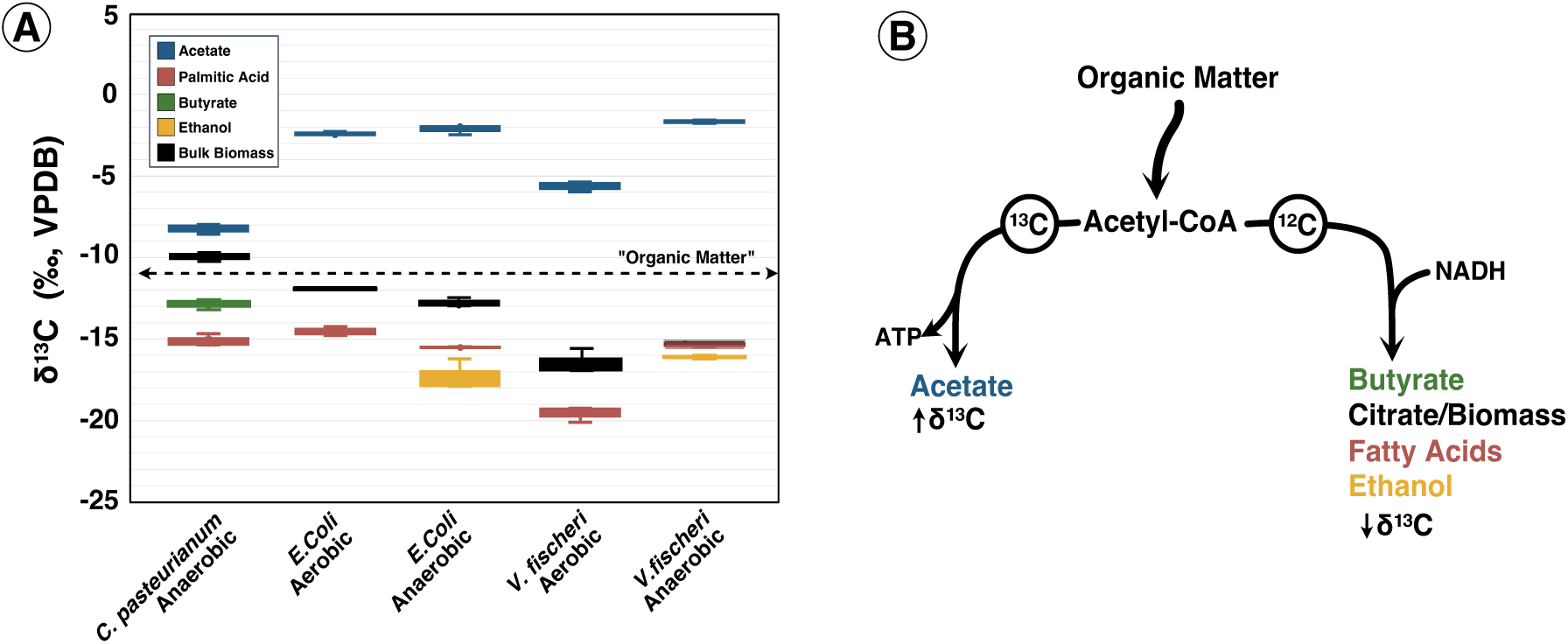
(A) Comparison of experimental results obtained from three fermenting organisms and five total conditions, includinga aerobic and anaerobic growth of *E. coli* and *V. fischeri* . Dotted line represents glucose molecular-average *δ*^13^*C* value. Box plots represent biological replicates. The carbon isotope composition of metabolic products that have acetyl-CoA as a precursor are shown to demonstrate that isotopic fractionations must occur at this node of metabolism to achieve the experimental results. (B) The schematic of carbon isotope flow at the acetyl-CoA node which creates ^13^*C*-depleted biomass, lipids, ethanol and butyrate and acetate ^13^*C*-enriched. Labels on the arrows denote high magnitdue KIEs (*<*0.99)of enzymes to butyrate, citrate, fatty acids, and ethanol and low magnitude KIEs of enzymes to acetate (∼1).

**Figure 4:**
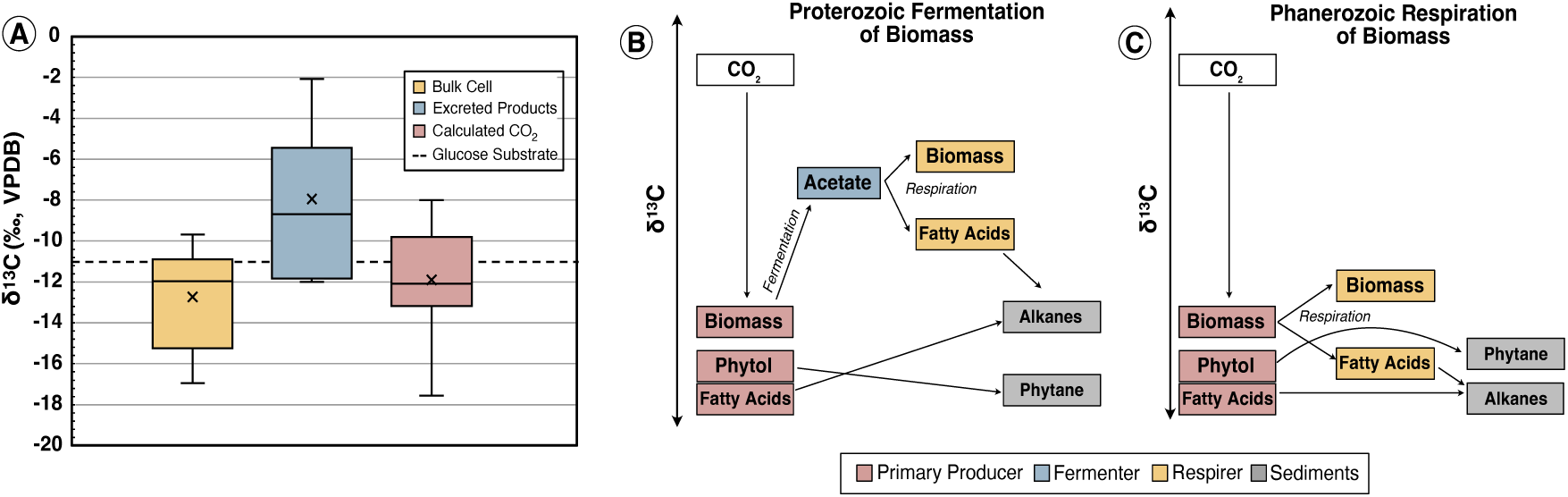
(A) Mass balance of carbon isotopes in different pools from all of the cultures in this study. Compound-specific *δ*^13^*C* values of fermentation products were averaged, weighting for carbon number and concentration. Carbon isotope compositions of excreted CO_2_ were not measured. Instead, they were calculated assuming all of the glucose consumed was converted to biomass, fermentation products and CO_2_ due to mass balance. Excreted fermentation products (blue) were on-average ^13^*C*-enriched compared to CO_2_ (red) and biomass (yellow). Panels B and C represent a schematic model of organic carbon flow under anoxic Proterozoic (B) and oxic Phanerozoic (C) pelagic water column conditions adapted from Logan et al. [1995] that leads to the observed isotopic pattern of lipid biomarkers in the rock record. When fermenters degrade primary produced cells (red), they excrete isotopically enriched fermentation products (blue), which are consumed by secondary degraders (yellow). These degraders contribute sedimentary n-alkanes with *δ*^13^*C* values (grey) exclusively in the Proterozoic when deep ocean anoxia was widespread. The isotopic inversion of phytane and n-alkanes may then be a signal of ancient microbial fermentation.

## Discussion

### Acetyl-CoA partitioning controls the ***δ*^13^*C*** patterns of excreted organic acids and lipid fatty acids

The consistent ^13^*C*-enrichment of acetate during fermentation in our results and others [Penning and Conrad, 2006] suggests a physiological mechanism shared by many fermentation pathways. The precursor to acetate, acetyl-CoA, plays a central role in bacterial metabolism as an entry point to lipid biosynthesis and the tricarboxylic acid (TCA) cycle. In fermenting bacteria, it is also used as an electron acceptor to generate products like butyrate, ethanol, and succinate. Despite having a shared precursor molecule, all four of these fermentation products and the C_16_ fatty acid palmitic acid had distinct *δ*^13^*C* values. As such, the reactions consuming acetyl-CoA must have variable carbon isotope KIEs. In our models, the KIE of enzymes that initiate the TCA cycle, lipid biosynthesis, ethanol production and butyrate synthesis all have strong KIEs which leaves the acetyl-CoA pool ^13^*C*-enriched (Table 1, Figure 3). Acetate inherits this ^13^*C*-enriched signature, resulting in high *δ*^13^*C* values of acetate excreted by three of the four bacteria. Meanwhile, fatty acids, ethanol and butyrate were ^13^*C*-depleted in those same organisms.

In fermenting cells, the acetyl-CoA node balances ATP generation and redox control. Synthesizing acetate is a significant energetic boon for fermenters because it generates ATP. However, this pathway consumes acetyl-CoA molecules that could otherwise be used as an anabolic precursor or as an electron acceptor to produce ethanol, butyrate, and succinate. Synthesizing these more reduced products regenerates NAD+ and other electron carriers that are required to catabolize more organic substrate. Depending on the organism’s genotype, its carbon substrate and its ambient environmental conditions, the branching ratio at acetyl-CoA will change. However, we predict that since the enzymes consuming acetyl-CoA induce *>*10‰ isotope effects, the residual *δ*^13^*C* of acetyl-CoA will always be enriched relative to the organic substrate (Figure 3). This includes citrate synthase, the enzyme that sends acetyl-CoA to the TCA cycle where amino acids and other cell-building precursors are synthesized. Indeed, even when *V. fischeri* and *E. coli* are grown under aerobic conditions, they express an overflow metabolism, known as aerobic fermentation [Wolfe, 2005, Dunn, 2012]. The acetate excreted under these conditions was enriched relative to glucose, owing to the KIE on citrate synthase, which likely shuttled most acetyl-CoA to the TCA cycle. In theory, as more acetyl-CoA is partitioned toward acetate and away from other acetyl-CoA pathways, acetate will approach the *δ*^13^*C* value of the fermentation substrate but will not become more depleted. We contend that this will hold true when other carbon substrates (i.e. amino acids, polysaccharides) are fermented, because there will still be a required balance of ATP generation, redox homeostasis and anabolism, all of which are controlled at the acetyl-CoA node. Thus, fermenting bacteria should consistently excrete acetate that is ^13^*C*-enriched relative to the consumed carbon substrates.

The preceding discussion warrants a reinterpretation of the carbon isotope signature found in bacterial fatty acids (e.g. palmitic acid), because acetyl-CoA is the building block of lipid biosynthesis. For decades, it has been assumed that fatty acids incorporate the carbon isotope composition of acetyl-CoA with minor fractionations imposed during lipid elongation [Hayes, 2001]. Acetyl-CoA was thought to inherit ^13^*C*-depleted carbon due to KIEs on the decarboxylation of pyruvate [Monson and Hayes, 1980, DeNiro and Epstein, 1977, Melzer and O’Leary, 1987]. However, our results demonstrate a consistent offset between acetate and palmitic acid *δ*^13^*C* values (Figure 3). In bacteria, these molecules are generated from the same pool of acetyl-CoA. Thus, the reaction from acetyl-CoA to malonyl-CoA, known as acetyl-CoA carboxylase, must have a strong isotope effect. Otherwise, fatty acids would inherit the same ^13^*C*-enriched carbon as acetate. The best fit across the four organisms is a C2 carbon isotope effect of 27‰. During *in vitro* studies of pyruvate carboxylase, which has an analogous reaction mechanism, the KIE on the methyl-site carbon isotope effect was 22‰, similar to our model results [O’Keefe and Knowles, 1986]. We therefore propose an alternative hypothesis in which acetyl-CoA polymerization is responsible for the ^13^*C*-depletion of fatty acid lipids in heterotrophic bacteria and possibly other organisms.

Shifting focus to acetyl-CoA carboxylase has important implications for the predicted intramolecular carbon isotope distribution in lipids. Its primary isotope effect putatively occurs on the methyl (C2) site of acetyl-CoA based on its reaction mechanism, creating an isotopic depletion at the odd carbon positions of fatty acids [Julien et al., 2022, O’Keefe and Knowles, 1986]. The site-specific distributions of carbon isotope compositions within fatty acids would then have an oscillating, see-saw pattern of relative isotopic enrichments and depletions along the alkyl chain at even and odd carbon positions respectively. However, this pattern would be inverted relative to empirical results in *E. coli* [Monson and Hayes, 1980] where the odd-numbered C9 carbon site was isotopically enriched relative to C10. Notably, the two sites measured in the previous study were part of an unsaturated carbon bond in the fatty acid molecule. It is possible that the isotopic patterns in these carbon atoms, with sp^2^ hybridizations do not represent — and should not be extrapolated to — the entire, saturated molecule. Consistent with our hypothesis, more recent data from saturated plant fatty acids find ^13^*C*-enrichment at odd-numbered carbon sites [Julien et al., 2022]. The nature of intramolecular carbon isotope patterns in lipids is an auspicious topic in geobiology for fingerprinting different forms of life throughout Earth history and astrobiology for identifying biotic molecules in extraterrestrial sample. As analytical techniques advance and more data becomes available, bioisotopic models like those developed here provide a quantitative perspective to interpret results and test hypotheses.

### Anaerobic heterotrophy may impose a trophic isotope fractionation

The small molecules produced by fermenting bacteria feed secondary heterotrophs like sulfate reducers. Our study demonstrates that these products are on average ^13^*C*-enriched relative to the starting substrate with individual organic acids inheriting even larger fractionations (Figure 4A). Metabolic models suggest that these fractionations are not specific to individual taxa or certain organic substrates. Rather, they derive from metabolic branch points that are widespread among fermentation pathways. While further experiments should test this hypothesis, the fractionations from fermentation likely persist across diverse bacteria and are relevant to environmental systems. In these systems, secondary degraders would inherit the isotopically enriched signature of their carbon source, imposing a trophic fractionation between local organic matter and secondary degraders. The magnitude and direction of this trophic fractionation would depend on which fermentation pathways are used and which fermentation products are preferentially consumed by terminal degraders. However, such trophic fractionations would only persist under anoxic conditions where fermentation is active. These ideas have implications in the search for a biosignature of fermentation in the Precambrian carbon isotope record.

### Molecular fossils elucidated the role of fermentation on the ancient Earth

The record of organic carbon burial within marine sedimentary rocks extends through the Archean era and is an important repository for studying ancient life on Earth. These deposits have primarily been interpreted as a signal of autotrophy, with variable *δ*^13^*C* values representing different proportions of carbon fixation pathways. Here we demonstrate that fermentation imposes its own isotopic fractionations that could be imprinted on the rock record, particularly in the Precambrian. In the following section, we investigate one potential signature of this metabolism that is preserved in residual sedimentary biomarkers.

A common feature of ancient sedimentary deposits is the isotopic inversion of alkane and phytane biomarkers across the Proterozoic-Phanerozoic boundary. This pattern is described as an “inversion” because it diverges from the *in vivo* isotopic ordering of biomolecules in algae and cyanobacteria. Modern phototrophs synthesize isoprenoids like phytol that are ^13^*C*-enriched relative to fatty acids. [Hayes, 2001]. These molecules are preserved as phytane and alkane biomarkers, respectively, and they retain their isotopic ordering (*δ*^13^*C _alkanes_ <δ*^13^*C _phytane_*) in Phanerozoic rocks. However, in sedimentary rocks of the Proterozoic, that ordering is consistently reversed (*δ*^13^*C _alkanes_ >δ*^13^*C _phytane_*). The original hypothesis for this inversion was an intense heterotrophic reworking of organic matter during the Precambrian [Logan et al., 1995]. Since heterotrophs synthesize fatty acids but do not produce phytol, their contribution to the sedimentary alkane pool could change the carbon isotope composition of alkanes relative to phytane. However, the trophic enrichment of carbon isotopes from primary producers to heterotrophs under aerobic conditions is not sufficient (∼1‰) to cause the observed isotopic inversion [Close et al., 2011].

The anoxic conditions of Precambrian ocean provides a unique scenario under which fermentation played a much larger role in recycling organic matter than in the oxic oceans of the Phanerozoic. As sinking particles were fermented, the trophic enrichment from organic matter to secondary anaerobic heterotrophs could have been up to 8‰ (Figure 4B). Such trophic enrichments have been observed in anoxic microbial mats analogous to Proterozoic conditions [Gonzalez-Nayeck et al., 2023]. As lipids derived from secondary heterotrophs slowly replaced those from phytoplankton, the total fatty acid pool would inherit the same isotopic enrichment, pushing its *δ*^13^*C* value higher than syngenetic phytol and inverting the isotopic ordering of sedimentary biomarkers. In the Phanerozoic, ocean basins became oxygenated and aerobic respiration replaced fermentation, degrading the majority of primary produced organic matter before it could reach anoxic sediments to be fermented. The trophic enrichment would therefore disappear in the sedimentary rocks of the Phanerozoic. Our hypothesis also provides a self-consistent explanation for the reappearance of this isotopic inversion during Phanerozoic marine anoxic events, which has been observed in sediments from the Permian [Grice et al., 2005, Nabbefeld et al., 2010], Ordovician [Pancost et al., 1999], and Triassic [Ajuaba et al., 2022, Fox et al., 2020] mass extinctions. In modern ocean sediments, we predict that the signal from fermentation could be seen in taxonomically specific fatty acids like the i17:1Δ9 biomarker associated with sulfate reducing bacteria that consume ^13^*C*-enriched fermentation products, though not in the total fatty acid pool, which contains many lipids from aerobic heterotrophs and terrigenous material.

Our study provides the first evidence that fermenting bacteria may have had an established role in the Precambrian marine carbon cycle, opening new questions about how this metabolic niche influenced other important biogeochemical processes such as nutrient recycling and carbon burial efficiency. The *δ*^13^*C* signature in ubiquitous lipid biomarkers could serve as a proxy to reconstruct the global importance of fermentation throughout Earth history. However, to fully elucidate the magnitude of the proposed fermentation ^13^*C*-fractionation on lipid biomarkers, anoxic degradation experiments need to be performed on natural organic matter. Future work should focus on developing such model systems. Metabolic bioisotopic models like those presented here provide a quantitative framework through which to predict and interpret data generated in experimental systems, in analogous modern environments, and in the rock record.

## Conclusions

Fermentation plays a crucial role in the Earth’s carbon cycle. Given its widespread importance across diverse ecosystems, tools to understand the activity and metabolic strategies of fermenting microorganisms are needed, both in modern environments and in the rock record. We demonstrate that fermenting bacteria growing on a simple sugar can express a large range of carbon isotope fractionations that are inherited by their excreted products. Using a combination of flux-balance analysis and bioisotopic modeling, we identified the enzymes consuming acetyl-CoA as the major drivers of the observed isotopic fractionations. Taking into account the physiological constraints on fermenting bacteria, we predict that the ^13^*C*-enrichment of acetate will be robust in the environment.We conclude that fermentation may impose a trophic isotope fractionation of up to 8‰ between organic matter (^13^*C*-depleted) and heterotrophic biomass (^13^*C*-enriched). Looking to the geologic past, such a trophic fractionation may explain the inverted isotopic ordering of isoprenoid and n-alkane biomarkers in the Precambrian. More broadly, our study demonstrates the utility of coupled metabolic models and isotopic measurements at their natural abundances. By identifying the mechanisms of isotope fractionations in microbial metabolism, we can evaluate the robustness of these signatures in the environment, generalize their application beyond pure cultures, and make predictions about how those signals will be preserved in the rock record.

## Materials and Methods

### Culturing Conditions

*Escherichia coli* (MG1655), *Clostridium pasteurianum*, *Zymomonas mobilis*, and *Vibrio fischeri* (MJ11) were grown on minimal, defined media with glucose as a sole carbon source. Other than *V. fischeri*, phosphate buffer was used to maintain a pH of 7. For *V. fischeri*, MOPS was used as a buffer at pH 7.2. All media compositions can be found in Tables S1-S5. Cultures were initially spread on agar plates from glycerol stocks (frozen at -80C) in small, anoxic chambers. Since all organisms are facultative or spore-forming, plates were removed from anoxia to streak single colonies. Colonies were inoculated into rich-media containing 5 g/L yeast extract and the media components listed in Table S1 - S5 using balch tubes sealed with a butyl rubber stopper and tin crimp foil. These were sparged with nitrogen through a needle for 5 minutes. After growth, these were transferred to pre-flushed Balch-type tubes containing 5 mL of minimal media and glucose. These pure cultures were passaged 2-3 times before inoculation into triplicate, 1L culture bottle containing 100mL of minimal media and glucose (1-2% transfer volume) that had been flushed with nitrogen for 20 minutes. The glucose fed to these cultures was previously characterized for its site-specific *δ*^13^*C* values using methods from Gilbert et al. [2009]. Throughout the growth, optical density (OD) measurements at 600 nm were made by extracting 1 mL of sample through a syringe into a plastic cuvette and rapidly measuring the density on a spectrophotometer. Culture purity was confirmed via microscopy.

### Organic acid quantification

At each growth timepoint, 0.5 mL of sample was removed, syringe filtered with a 0.22 *µ*m filter and stored at -20C. These samples were diluted 2-fold into 8mN sulfuric acid and injected (10 *µ*L) onto a high-performance liquid chromatograph (HPLC) with a refractive index detector (RID) equipped with an autosampler. Sugars, organic acids and alcohols were separated on a Aminex-87H column with an isocratic method using 8mN sulfuric acid at 0.6 mL/min. The concentrations of glucose, succinate, acetate, ethanol, formate, and lactate were simultaneous quantified by running external standard curves (0.1-50 mM) of these analytes. In the case of *V. fischeri*, ethanol and MOPS co-eluted, precluding HPLC quantification of ethanol in those samples. To circumvent this issue, we measured ethanol on a gas chromatography (GC) instrument with flame ionization detection (FID). Here, samples were diluted two-fold into high-purity MilliQ water and injected (0.3 *µ*L) onto a ZB-WaxPlus column with a constant 120*^◦^*C injector temperature (split ratio = 10). Concentrations were quantified with an external standard curve. Three water rinses of the autosampler syringe before and after injections minimized sample carry over.

### Organic acid isotopic analyses

Compound-specific isotopic analyses of sugars, volatile fatty acids and alcohols were performed on an isotope ratio monitoring liquid chromatography/mass spectrometry system (irm-LC/MS) at the University of Bremen. The analysis involved separation of compounds by high performance liquid chromatography (Thermo Scientific^TM^ Dionex UltiMate 3000 HPLC) combined with chemical oxidation of the effluent using the Thermo Scientific^TM^ LC IsoLink^TM^ interface and subsequent online transfer of the resulting CO_2_ into an irm-MS (ThermoFinnigan Delta Plus XP), using helium as a carrier gas. The method was isocratic and isothermal. All solutions were aqueous, freshly prepared and degassed under vacuum in an ultrasonic bath (15 min at 40°C) in order to remove CO_2_. The HPLC system was equipped with a VA 300/7.8 Nucleogel Sugar 810 H column (300 mm length; 7.8 mm i.d.) and a guard column (CC30/4 Nucleogel Sugar 810H; 30 mm length) from Macherey-Nagel, which were kept at 45°C. As a mobile phase, 5 mM phosphoric acid (prepared from 270 *µ*L of 85% H_3_PO_4_ in 1 L of MilliQ water) was used with a flow rate of 300 *µ*L min^-1^. The oxidation reactor temperature of the LC IsoLink^TM^ interface was set to 99.9°C. Inside the interface, the eluent of the HPLC was mixed with oxidation reagent prepared from two solutions, one being 283 mM sodium peroxodisulfate solution (prepared from 27 g Na_2_S_2_O_8_ in 400 mL of MilliQ water) and the other 1.4 M phosphoric acid (prepared from 30 ml of 85% H_3_PO_4_ in 400 ml of MilliQ water). The reagents were pumped into the interface with equal flow rates of 50 *µ*L min^-1^, adjusted to generate an oxidative power that yielded 11.5 V for oxygen measured as m/z 32 on cup number 2 (resistor of 3·10^8^ *ω*). Isotope-ratio-monitoring was conducted on a ThermoFinnigan Delta Plus XP, to which the interface was connected with helium as carrier gas. Samples were injected using a temperature-controlled autosampler (set to 5°C), and a sample injection volume of 100 *µ*l. Analysis were repeated with diluted samples when peak areas of one or several target compounds exceeded 500 Vs. In house isotopic standards of known *δ*^13^*C* value were run at the beginning of the analytical sequence and in between every seven samples to ensure accuracy throughout. Precision ranged from 0.2‰ for ethanol, glucose and succinate to 0.7‰ for lactate and propionate. In samples, isotopic analysis of lactate suffered from co-elution with succinate when the latter was present in high concentrations. *δ*^13^*C* values are reported by comparison to Vienna Peedee Belemnite (VPDB).

### Lipid isotopic analyses

Fatty acid carbon isotope compositions were determined via gas chromatograph isotope ratio mass spectrometer method. Fatty acids were first transesterified to fatty acid methyl esters (FAMEs). To do so, 1mL of GC-purity hexanes and 2 mL of a 20:1 mixture of anhydrous methanol and acetyl chloride were added to 5-10 mg of freeze-dried and crushed bacterial biomass. This mixture was heated to 100*^◦^*C for 10 minutes. After allowing the solution to cool to room temperature, a liquid-liquid extraction with hexanes and water was performed. The organic phase was extracted and dried using a sodium sulfate column.

The FAME profile of each organism was determined using a Thermo Scientific Trace 1300 ISQ with separation on a 30-m × 0.25-mm capillary column (ZB-5 ms, 1 *µ*m film thickness; Zebron). The *δ*^13^*C* values of FAMEs were measured by a gas chromatograph coupled to an isotope-ratio mass spectrometer (Thermo-Scientific Delta+XP) using a combustion interface. Chromatographic separation was carried out with the same column. Identical oven temperature ramp settings were used for GC/MS analysis, so that peaks could be identified on the GC-IRMS by relative retention time. For each sample, triplicate measurements were performed to determine *δ*^13^*C* values. The average *δ*^13^*C* values were corrected for the added methyl C in the derivatization. An internal reference standard of CO_2_ gas bracketed the chromatographic FAME peaks to determine their isotope composition on the VPDB scale. Biological reproducibility was consistently *<*0.5‰.

### Flux-balance metabolic models

To construct the metabolic models for each organism, flux balance analysis was used. Previous investigations of fermentation pathways utilized by each bacterium were leveraged[Dunn, 2012, Jacobson et al., 2019, Sprenger, 1996, Clark, 1989, Heyndrickx et al., 1986]. From these studies, we created a set of four master flux maps to represent carbon flow in the organisms (Figure S4). The internal fluxes were constrained by the glucose uptake flux, fermentation product excretion fluxes and anabolic fluxes to key biosynthetic pathways including amino acids, fatty acids, and nucleic acids. We determined the latter rates by assuming a cellular molecular composition (moles/cell) similar to that of *E. coli* as determined by Neidhartd (1982). By normalizing to the growth rate and cell densities, these were converted to flux terms (moles/g dry cells/hr). The excretion and uptake rates were constrained by plotting the concentration of these substrates and products versus the cell density for all biological replicates of each organism. The fitted linear slope of this plot was taken as the excretion and uptake rates (moles/g dry cells/hr).

A system of equations to describe the internal fluxes of each cellular metabolic reactions was derived from the known stoichiometries of these reactions (Supplementary Information). Each system of equations for the different organisms is fully constrained by the empirical uptake, excretion, and anabolic fluxes (Supplementary Figure S4 and S5, Table S6). For visualization purposes in Figure 1, the reaction fluxes were normalized to the glucose uptake rate and then multiplied by 50. They represent moles of substrate, so when all export fluxes are multiplied by their respective carbon numbers, the sum is 300, matching the uptake flux (50 × 6 carbons in glucose).

### QIRN Models

A detailed description of QIRN’s methods for modelling isotope fractionations within reaction networks is published elsewhere [Mueller et al., 2022]. Briefly, we constructed the flux maps of fermentative metabolism that are abbreviated in Figure 1 and Figure S5. We assigned kinetic isotope effects to those enzymes listed in Table 1 at their designated atomic sites. KIEs were also assigned as 0.987 and 0.99 on the C1 and C2 sites, respectively, of pyruvate cleavage into acetyl-CoA and formate (pyruvate formate lyase, in *E. coli* and *V. fischeri*) and as 0.987 and 0.99 on C1 and C2, respectively of pyruvate cleavage into acetyl-CoA and CO2 (pyruvate ferrodoxin oxidoreductase, in *C. pasteurianum*). However, these enzymatic KIEs had no effect on the RMSE of the model-data comparisons as discussed in the main text. As such they were not included in Table 1.

It was assumed that these KIE’s all were the same in the four organisms. The *δ*^13^*C* values of all the organisms’ fermentation products(n=17) could then be used as constraints on the KIEs. QIRN looped through a parameter space from 0.96 to 1 for each of these enzymatic KIEs, testing the residuals between model predictions and data for each simulation. In these tests, two enzymatic KIEs were varied simultaneously while holding all others constant (Figure S5). The total parameter space was not searched as this would create intractably long computation times (∼months). All QIRN models were run for 2400 timesteps (dimensionless) which gave steady state isotope compositions for all end products. Rather than synthesizing a C_18_ fatty acid, which has over 250,000 isotopologues, we calculated its theoretical *δ*^13^*C* value from the steady-state site-specific isotope compositions of its precursors, acetyl-CoA and malonlyl-CoA. Given the unidirectional nature of fatty acid biosynthesis, we assumed there was negligible fractionation from these monomers to the final lipid product. Palmitic acid is synthesized from an aceytl-CoA starting block and seven molecules of malonyl-CoA subsequently elongated off of this acetyl-CoA.

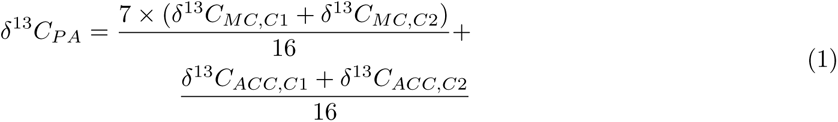

Thus its *δ*^13^*C* value(*δ*^13^*C_P A_*) is a function of the site-specific *δ*^13^*C* values of malonyl-CoA’s C1 and C2 sites (*δ*^13^*C_MC,C_*_1_ and *δ*^13^*C_MC,C_*_2_) and the site-specific *δ*^13^*C* values of the C1 and C2 sites in acteyl-CoA (*δ*^13^*C_ACC,C_*_1_ and *δ*^13^*C_ACC,C_*_2_). We divided these by 16, the number of carbons in palmitic acid, to derive the molecular average *δ*^13^*C* value.

## Acknowledgements

The authors gratefully acknowledge Professor G’erald Remaud for intramolecular isotopic analysis of glucose stocks. We thank Professor James B McKinlay and Professor Ned Ruby for offering strains of *Z. mobilis* and *V. fischeri*, respectively. We thank Victoria Orphan for use of her laboratory facilities and Ludmilla Aristilde and Roger Summons for helpful discussions in the interpretation of the data. We are also grateful for the contributions of Jenny Wendt from MARUM (University of Bremen) and Daniel Felsmann from Thermo Fisher Scientific for their competent and dedicated support in operating the isotope ratio monitoring liquid chromatography/mass spectrometry system. Funding for this work came from an NSF Graduate Research Fellowship DGE-1745301 (to E.P.M.), a European Association of Organic Geochemistry Research Award (to E.P.M.) and the NASA Astrobiology Institute grant # 80NSSC18M0094 (to A.L.S.). This work was also supported by the Deutsche Forschungsgemeinschaft through the Cluster of Excellence “The Ocean Floor – Earth’s Uncharted Interface” (project 390741603) (to V.B.H. and K.-U.H.).

## Author Contributions

E.P.M conceived of the study, cultured the microorgnasisms, and wrote the manuscript. E.P.M and V.B.H. performed isotopic and chemical analyses. E.P.M., A.L.S., K.H., and V.B.H. interpreted the data. K.H., A.L.S., and J.L. contributed analytical and biological laboratory facilities. All authors contributed to the editing of the manuscript.

## Notes

### Competing Interest Statement

The authors have declared no competing interest.

