## Supplementary Materials for "Metabolic controls on carbon isotope fractionations during bacterial fermentation"

**Supplementary Information**
Metabolic controls on the carbon isotope fractionation of bacterial fermentation
Mueller *et al.*

### Microbial Growth Conditions and Data

*Escherichia coli*, *Vibrio fischeri*, *Zymomonas mobilis*, and *Clostridium pasteurianum*were grown under fermenting conditions with glucose as a sole carbon source. The media conditions for each organism are below in Tables S1 – S4. All organisms were grown on a glucose stock with a known intramolecular isotopic structure (Table S5).

**Table S1***: Z. mobilis*

| Species | Concentration (g/L) |
| --- | --- |
| $MgCl_2*6H_2O$ | 0.4 |
| NaCl | 1 |
| $CaCl_2*6H_2O$ | 0.1 |
| KCl | 0.5 |
| $CaCl_2*2H_2O$ | 0.07 |
| $NH_4Cl$ | 0.53 |
| $Na_2SO_4$ | 0.04 |
| $K_2HPO_4$ | 0.17 |
| $KH_2PO_4$ | 0 |

**Table S3:** *V. fischeri*

| Species | Concentration (g/L) |
| --- | --- |
| $MgCl_2*6H_2O$ | 8.73 |
| NaCl | 17.46 |
| $CaCl_2*6H_2O$ | 0.13 |
| KCl | 0.44 |
| $CaCl_2*2H_2O$ | 0.09 |
| $NH_4Cl$ | 0.8 |
| $Na_2SO_4$ | 3.55 |
| $K_2HPO_4$ | 0.17 |
| $KH_2PO_4$ | 0 |

**Table S2:** *C. Pasteurianum*

| Species | Concentration (g/L) |
| --- | --- |
| $Na_2HPO4*12H_2O$ | 2.2 |
| $KH_2PO_4$ | 5.97 |
| $(NH_4)_2SO_4$ | 6 |
| $MgSO_4*7H_2O$ | 0.1 |
| $CaCl_2$ | 0.01 |
| $Na_3Citrate*2H_2O$ | 0.33 |

**Table S4:** *E. coli*

| Species | Concentration (g/L) |
| --- | --- |
| $Na_2HPO_4*7H_2O$ | 11.32 |
| $KH_2PO_4$ | 3 |
| NaCl | 0.5 |
| $(NH_4)_2SO_4$ | 2.5 |
| $MgSO_4$ | 0.120 |
| $CaCl_2$ | 0.011 |

**Table S5:** Glucose intramolecular $\delta^{13}C$ composition

| Carbon Site | Isotope Composition (‰) |
| --- | --- |
| 1 | -6 |
| 2 | -7.8 |
| 3 | -8.5 |
| 4 | -14 |
| 5 | -14.9 |
| 6 | -14.7 |

Growth was measured as optical density (OD) for each organism from lag phase to mid-exponential phase, at which point the cultures were rapidly collected for biomass. Growth rates were calculated from the exponential portion of each growth curve.

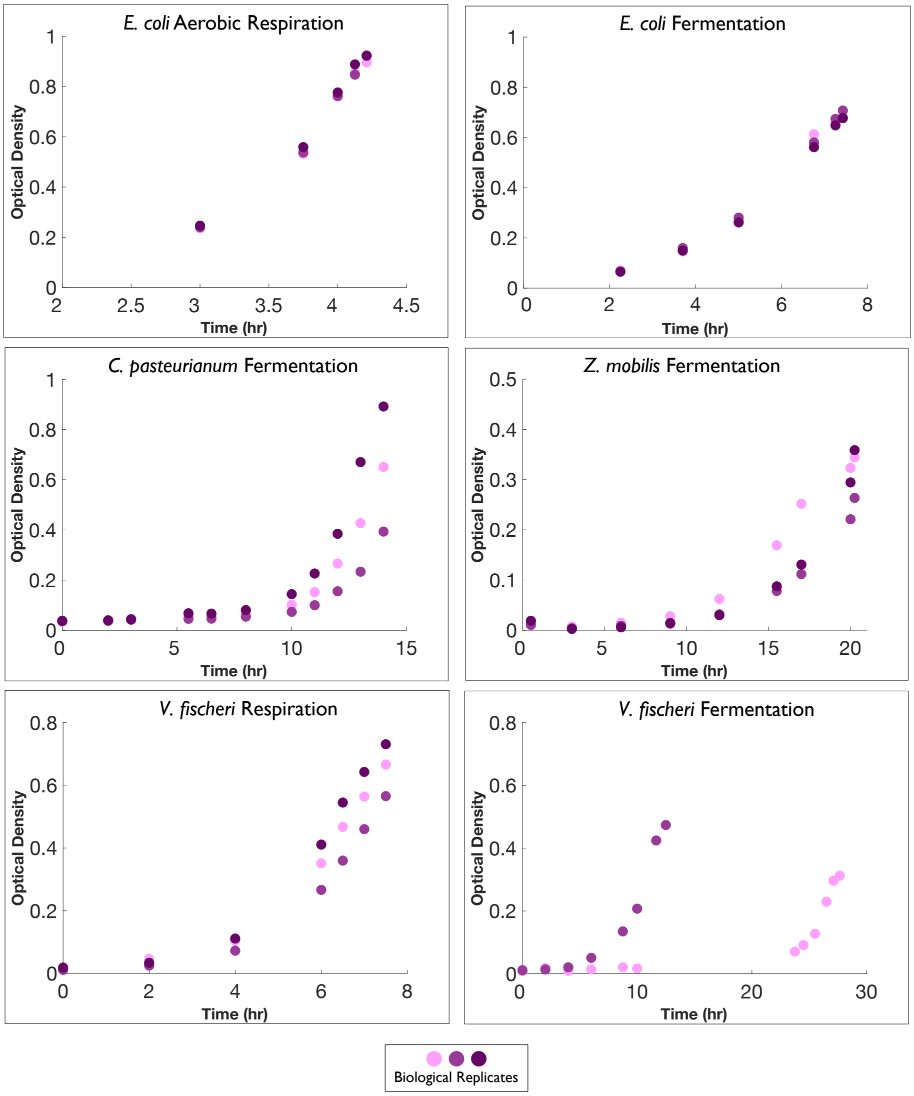


Fig. S1: Growth curves of the four model organisms, Escherichia coli, *Vibrio fischeri*, *Zymomonas mobilis*, and *Clostridium Pasteurianum*. *E. coli* and *V. fischeri* were grown aerobically and anaerobically.

Organic acid profiles were measured via high-performance liquid chromatography (HPLC) at the same time that cells were collected. Biological triplicates demonstrate high biological reproducibility.

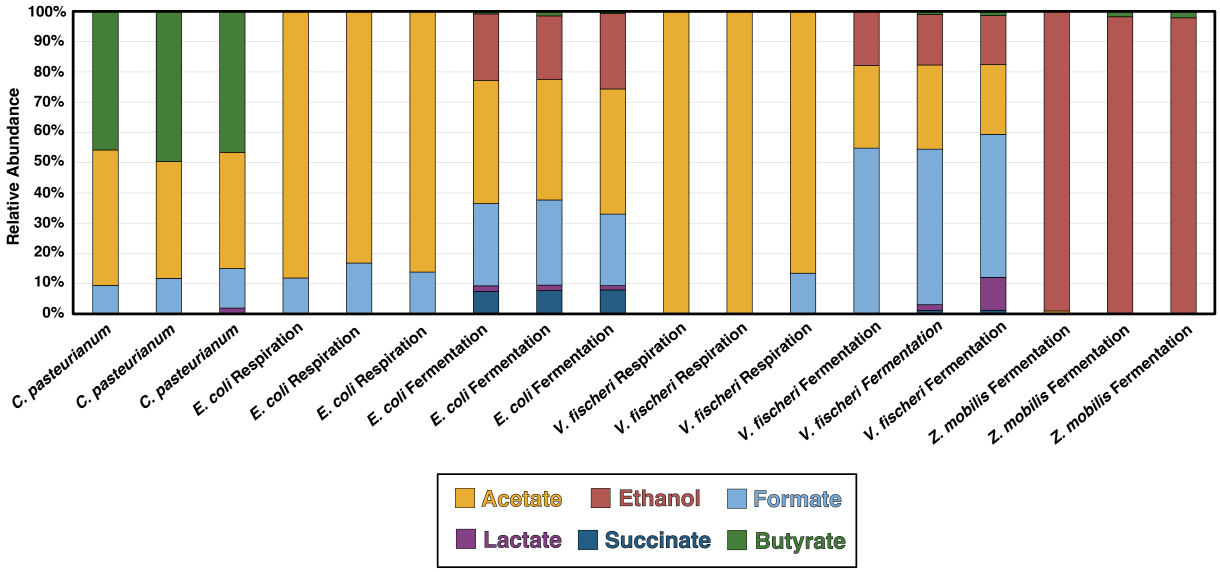


Fig. S2: Relative abundance of fermentation product concentrations in mid-exponential phase at the point of biomass collection.

### Building metabolic models

To constrain the metabolic controls on the diverse isotopic fractionations associated with microbial fermentation that we measured here, metabolic models were built via flux balance analysis (FBA). To perform FBA, the substrate uptake flux must be balanced by product excretion (organic acids, alcohols and gases) and anabolism fluxes (DNA, proteins, and lipids). Fermentation products and glucose concentrations were measured throughout the growth to quantify excretion and uptake rates, respectively. Cell density (g/L) was quantified by multiplying optical density (OD 600) by a constant (0.56). In Figure S3, data from biological replicates were collated into a single data set to constrain the slopes in Figure S3.

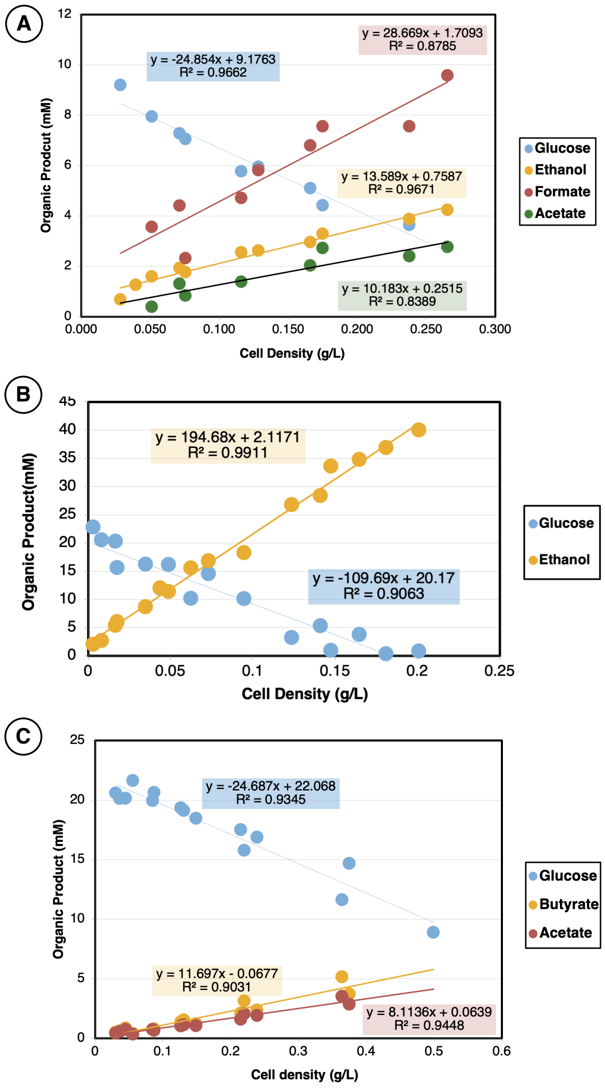


Fig. S3: Relationships between time-varying cell densities (grams cell dry weight per liter), glucose concentration, and fermentation product concentrations. The slopes of these correlations are the zeroth-order fluxes used to constrain the flux balance analysis in terms of moles per gram cell dry weight per hour. A.) *V. fischeri*, B.) *Z. mobilis*, C.) *C. pasteurianum*

The anabolic fluxes were determined using the cellular composition of *E. coli* (in moles/gram cell dry weight) as determined in Neidhart (1987) (Table S6). These values were then multiplied by the growth rate ($hr^{-1}$) to yield a flux term (moles/L/hr).[1] For protein synthesis, amino acids were categorized by family and fluxes from central metabolism were assigned by summing the fluxes of individual amino acids in those families (Table S6). Similarly, the total flux to DNA and RNA was estimated by the sum of the fluxes to individual nucleotides.

With the excretion, uptake, and anabolic fluxes determined, the internal fluxes were constrained using a system of equations that describes the fermentation pathways in Figure S4. This system of equations only coarsely parameterized metabolic pathways (e.g., glyceraldehyde-3-phopshate to 3-phosphoglycerate), whereas the QIRN models utilized to create Figure 2 in the text included all intermediary steps.(e.g. glyceraldehyde-3-phopshate to 1,3-bisphosphoglycerate to 3-phosphoglycerate). The fluxes of these intermediary steps are equivalent and would be redundant in Figures 1 and S4.

This metabolic map is described by a system of equations incorporating known stoichiometry of these enzymatic reactions. Equations shifted slightly for *Z. mobilis*, which utilizes the Entner-Dodouroff pathway and creates ethanol directly from pyruvate rather than from acetyl-CoA. The fluxes determined using the system of equations were then implemented into the larger QIRN models, which are attached as supplementary files. In QIRN, the starting concentration of glucose was set to 0.5 (arbitrary units), so the forward rate constants for the reactions upstream of aldolase (F6P to DHAP and GAP) in the Network Builder were doubled to achieve the fluxes found in Figure 1. Only one

| Table S6 | | | | |
|---|---|---|---|---|
| **Amino Acids** | **umol/g cell** | | **Nucleotides** | **umol/g cell** |
| ALA | 488 | | AMP | 165 |
| ARG | 281 | | GMP | 203 |
| ASX | 458 | | CMP | 126 |
| CYS | 87 | | UMP | 136 |
| GLX | 500 | | DAMP | 24.6 |
| GLY | 582 | | DGMP | 25.4 |
| HIS | 90 | | DCMP | 25.4 |
| ISO | 276 | | DTMP | 24.6 |
| LEU | 428 | | | |
| LYS | 326 | | **Lipids** | **umol/g cell** |
| MET | 146 | | Fatty Acids | 258 |
| PHE | 176 | | | |
| PRO | 210 | | **AA Families** | **AAs** |
| SER | 205 | | Pyruvate | ALA, VAL, ISO, LEU |
| THR | 241 | | 3PGA | GLY, SER |
| TRP | 54 | | OAA | ASX, ARG |
| TYR | 131 | | CIT | GLX,MET,PRO,THR |
| VAL | 402 | | R5P | HIS, PHE, TRP, TYR |

*All data from Neidhardt, 1987

anabolic pathway incorporated inorganic carbon into meatbolites, phosphoenolpyruvate (PEP) carboxylase (PEP + $CO_2$ to oxaloacetate). This $CO_2$ pool was assumed to come directly from metabolic reactions within the cell. Specifically, $CO_2$ from formate dehydrogenase was diverted to feed the PEP carboxylase reaction.

$$\phi_2 = \phi_G - \phi_{DNA} - \phi_{AA1} \tag{1}$$

$$\phi_3 = 2 \times \phi_2 \tag{2}$$

$$\phi_4 = \phi_3 - \phi_{AA2} \tag{3}$$

$$\phi_5 = \phi_4 - \phi_6 \tag{4}$$

$$\phi_7 = \phi_5 - \phi_{AA3} - \phi_L \tag{5}$$

$$\phi_8 = \phi_7 - \phi_{AC} - \phi_{FA} - \phi_E - 2 \times \phi_B \tag{6}$$

$$\phi_6 = \phi_8 + \phi_{AA4} + \phi_S \tag{7}$$

$$\phi_9 = \phi_7 - \phi_F \tag{8}$$

### Model Sensitivity Analysis

To evaluate the sensitivity of QIRN models to individual enzymes' KIEs, we varied two enzymatic KIEs independently and calculated the model-data residual as the root-mean square error with each simulation. In Figure S5 (left), carbon isotope data from fatty acids, acetate, ethanol and butyrate were used as constrains. In Figure S5 (right), formate and lactate carbon isotope compositions were used as constraints. To define the range of optimized KIE values (Table 1, in the main text), we set a limit of 3‰ for the RMSE which is three-fold higher than the analytical error of 1‰. These analyses demonstrated that acetyl-CoA carboxylase, alcohol dehydrogenase, and formate dehydrogenase can be constrained to varying extents. Acetyl-coenzyme A acetyltransferase is similarly constrained though not shown. Thus, while QIRN allows us to identify the enzymes that must have kinetic isotope effects, it does not uniquely constrain the exact isotope fractionation on those enzymes. Nor does QIRN identify which carbon site(s) the KIE is occurring on. These isotope effects are placed exclusively at carbon sites that are involved in the reactions (e.g. bond rearrangements, hybridization changes, bond cleavage, or condensation).

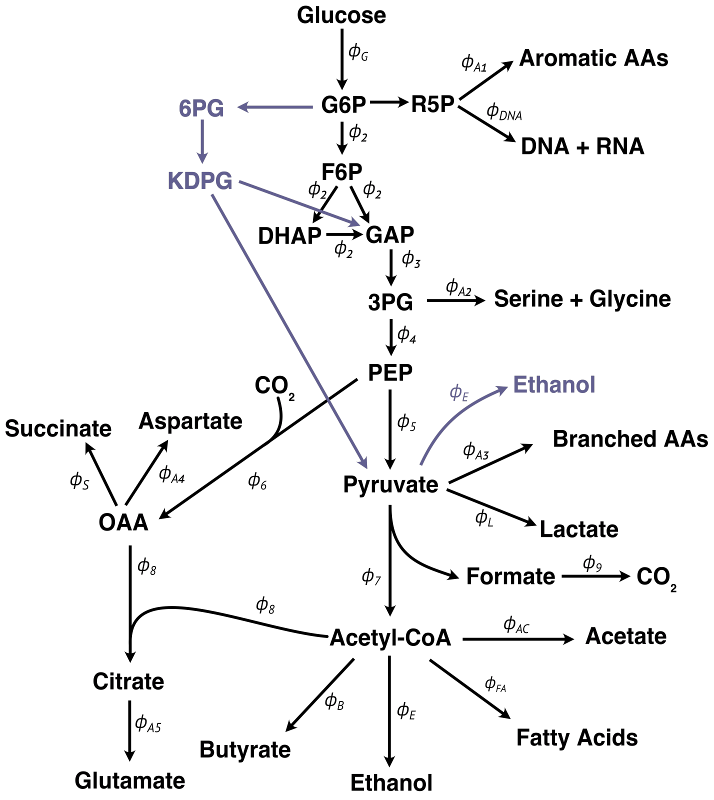


Fig. S4: Master metabolic map for the four organisms grown in this study. Fluxes ($\phi$) denoted with alphabetical subscripts represent anabolic and excretion fluxes that are estimated and empirically derived, respectively. Internal fluxes deduced from the system of equations below are denoted by numerical subscripts. Purple reactions denote those utilized by Z. mobilis but not the other organisms in this study. The reaction networks implemented into QIRN include the intermediates between major metabolic modes highlighted here (e.g. acetoacetyl-CoA, acetaldehyde, malonyl-CoA, etc.)

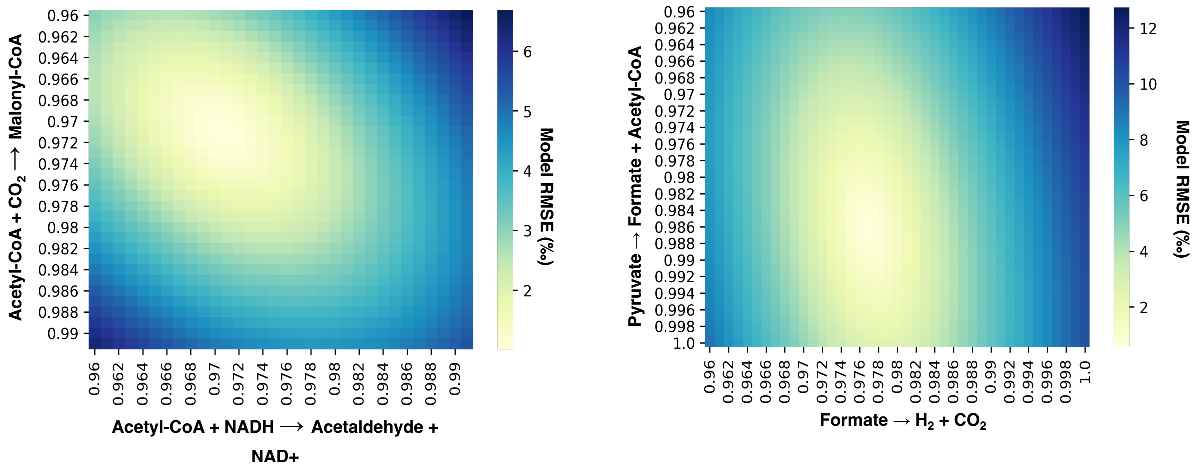


Fig. S5: Sensitivity analysis of four enzymes kinetic isotope effects. Color gradient indicates the model-data residuals calculated as root-mean-square-error (RMSE). Carbon site of KIE is indicated in parentheses next to the enzyme name.
